# BCG Vaccination at Birth Shapes the TCR Usage and Functional Profile of MR1T Cells at 9 Weeks of Age

**DOI:** 10.1101/2025.10.09.681353

**Authors:** Dylan Kain, GW McElfresh, Gwendolyn M. Swarbrick, Katherine Rott, Gregory Boggy, Gerhard Walzl, Nelita Du Plessis, Willem Hanekom, Elisa Nemes, Muki Shey, Thomas J. Scriba, Benjamin N. Bimber, David M. Lewinsohn, Deborah A. Lewinsohn

**Affiliations:** Division of Infectious Diseases, Department of Pediatrics, Oregon Health & Science University, Portland, OR, United States; Division of Pulmonary, Allergy and Critical Care Medicine, Oregon Health & Science University, Portland, OR, USA; Division of Infectious Diseases, Department of Medicine, University of Toronto, Toronto, ON, Canada; Oregon National Primate Research Center, OHSU, Beaverton, OR, USA; DSI-NRF Centre of Excellence for Biomedical Tuberculosis Research; South African Medical Research Council Centre for Tuberculosis Research; Division of Immunology, Department of Biomedical Sciences, Faculty of Medicine and Health Sciences, Stellenbosch University, Cape Town, South Africa; South African Tuberculosis Vaccine Initiative, Division of Immunology, Department of Pathology and Institute of Infectious Disease and Molecular Medicine, University of Cape Town, Cape Town, South Africa; Vaccine and Gene Therapy Institute, OHSU, Beaverton, OR, USA

**Keywords:** MR1, MAIT cells, MR1T cells, T cell development, Single-cell sequencing, Neonatal sepsis, Tuberculosis

## Abstract

Tuberculosis (TB) is the leading infectious disease killer worldwide and children suffer disproportionately. The ongoing burden of disease, despite widespread vaccination with BCG, highlights the need for novel vaccines. MR1-restricted T (MR1T) cells recognize small molecules, including microbial-derived molecules, presented by the monomorphic MHC class 1- related molecule (MR1). They have both “innate” effector capacity, allowing them to quickly respond to pathogens including *Mycobacterium tuberculosis* (Mtb), while also having adaptive features (effector memory cell surface phenotype and selective TCR usage). Feasibility of an MR1T cell-based vaccination remains unexplored and critical to this is whether or not MR1T cells possess the capacity for immunological memory. To begin to address this question, peripheral blood mononuclear cells (PBMC) were collected at 9-weeks of age from healthy term-infants in South Africa, who had either received BCG vaccination at birth (n=10) or who had BCG vaccination delayed (n=10). MR1/5-OP-RU tetramer positive cells were sorted using flow cytometry and single-cell RNA and TCR-seq was performed. *Ex-vivo* MR1T cells from vaccinated infants demonstrated increased expression of type I interferon response genes consistent with a cytokine mediated response to BCG vaccination. Using the TCR clustering algorithm TCRdist3, similar TCRs were grouped together, revealing a cluster significantly enriched in BCG-vaccinated infants. This cluster exhibited elevated expression of pro-inflammatory and cytotoxic genes, consistent with a recall response to prior vaccination and evidence of possible recognition of Mtb. This work provides the first step in addressing if MR1T cells demonstrate immunological memory, however, further work is needed to understand if these clonal expansions persist and possess the capacity for antigenic recall.

## Introduction

With 1.3 million deaths in 2022, tuberculosis (TB), caused by *Mycobacterium tuberculosis* (Mtb), remains the leading global infectious disease cause of death.^1^ Outcomes after exposure to Mtb vary widely however, with most individuals able to contain or clear infection, while some develop progressive disease.^2^ The immune system’s role in these divergent outcomes is underscored by the higher rates of severe disease in children.^3–5^ The only licensed vaccine for TB, bacille Calmette-Guérin (BCG) has not shown consistent benefit in adults,^6^ but has partial protection against severe disease in early life. Despite this, TB still causes approximately 200,000 deaths in children annually.^1^ Clarifying the basis of this incomplete protection is critical to advancing better vaccine strategies. In particular, understanding the role of T cells that recognize Mtb-infected cells, including non-conventional T cells like MR1-restricted T (MR1T) cells, may offer key insights into protective immunity.

MR1T cells are a type of donor-unrestricted T cells (DURTs), that recognize bacterial and fungal metabolites presented from the monomorphic MHC class 1-related molecule (MR1). While MR1T cells can be activated through their T-cell receptor (TCR), they can also be activated directly via cytokines.^7^ Mucosal associated invariant T (MAIT) cells are the canonical MR1T cell characterized by an invariant TCR α-chain (TRAV1.2-TRAJ33/20/12 in humans), paired with a variety of Vβ segments,^8–11^ and are generally CD8^+/-^CD4^-^CD161^++^CD26^+^.^12–14^ Using an MR1-tetramer, we and others have previously described MR1-restricted T cells that do not express TRAV1-2^15^ and therefore define MAIT cells as a subset of MR1T cells. In adults, TRAV1-2(-) MR1T cells are rare, but at birth they are the majority of MR1T cells.^16^ This balance rapidly shifts postnatally, and by 10 weeks, most MR1T cells resemble adult MAIT cells.^16^ This early maturation, faster than conventional T cells, suggests MR1T cells may contribute to early-life immune protection.^16–18^

MR1T cells are enriched in mucosal surfaces, including the lung, where they can represent over 10% of all T cells.^19^ They mediate rapid effector responses and contribute to defense against respiratory pathogens including *Klebsiella pneumoniae,*^20^ *Escherichia coli,*^21^ *Staphylococcus aureus,*^22^ *Streptococcus pneumoniae,*^23^ *Francisella tularensis,*^24^ and mycobacterial pathogens.^25^ Unlike conventional CD4/CD8 T cells, MR1T cells possess immediate effector function, and are able to kill invading pathogens much more rapidly. In humans, MR1T cells have been shown to be enriched in the lungs of those with active TB,^26^ and in those who are resisters to Mtb infection.^27^ Polymorphisms in MR1 have been associated with TB susceptibility,^28^ and MR1- deficient mice are more vulnerable to infection with *M. bovis*, but not Mtb.^25^ In non-human primates, depletion of MR1T and other non-conventional T cells worsens TB disease.^29^ These findings suggest MR1T cells are uniquely poised to provide rapid and early protection in the lung following aerosolized Mtb exposure – potentially critical in early life before the adaptive immune system fully matures.

Despite the potential role MR1T cells may play in defense against Mtb, the feasibility of MR1T based vaccination remains unexplored. MR1T cells are currently defined using a MR1/5-(2- oxopropylideneamino)-6-D-ribitylaminouracil (5-OP-RU)-tetramer, but despite all MR1T cells recognizing 5-OP-RU, there is clear evidence of antigen discrimination by MR1T cells,^30,31^ suggesting a biological plausibility for MR1T-based vaccination. It is not, however, currently known if these cells possess features of immunological memory, a critical component needed to determine how to utilize these cells for future vaccination.

Previous work has shown that after BCG vaccination at birth, there were no significant shifts in MR1T cell frequency,^32^ but there may remain shifts in TCR usage as a result of vaccination that could suggest immunological memory. To begin to address this question, we utilized biorepository samples from a previous study of South African infants who had either received BCG vaccination at birth (“vaccinated”) or who had BCG vaccination delayed until after peripheral blood mononuclear cells (PBMC) was collected at 9-weeks of age (“unvaccinated”).^32^ MR1T cells were sorted from these samples using a tetramer and herein we show that MR1T cells from vaccinated babies had significant upregulation of interferon stimulated genes (ISG), potentially acting in an antigen-independent, cytokine-driven manner to BCG. Furthermore, by using single-cell TCR-sequencing and the TCR clustering algorithm, TCRdist3,^33^ we show that vaccinated infants had a significant increase of one TCR cluster compared to unvaccinated infants. This TCR group had significant upregulation of pro-inflammatory genes, and downregulation of naïve genes suggestive of a mature effector phenotype. Moreover, we isolated an MR1T cell clone with a TCR that clustered into this enriched cluster, and we have previously shown that this clone recognizes Mtb.^31^ Overall this work suggests that BCG shapes MR1T cells both in a cytokine mediated manner but also suggests that there are antigen-specific changes mediated *in vivo* that persist 9-weeks after vaccination. This work provides the first step in addressing if MR1T cells demonstrate immunological memory, however, further studies are needed to understand if these clonal expansions persist and possess the capacity for antigenic recall.

## Results

### MR1T Cells Mature Quickly After Birth

To define the role of BCG vaccination on the development of human MR1T cells, single-cell gene RNA-sequencing (scRNA-seq) was performed on MR1/5-OP-RU-tetramer sorted MR1T cells from South African cord blood (CB, n=10), infants (n=20) at 9-weeks of age, or adult donors (n=5) (Figure 1A). Half of the infant donors had received BCG-vaccination at birth (n=10) and half had BCG vaccination delayed until after PBMC collection at 9-weeks of age (n=10). Representative flow gating strategy is shown in Supplemental Figure 1. Donor characteristics are summarized in Table 1. Cell surface protein expression was also examined through 5 oligo-conjugated antibodies. After filtering, there were 3,350 CB-derived, 3,400 unvaccinated infant-derived, 2,761 BCG-vaccinated infant and 13,535 adult-derived MR1T cells included in the analysis.

**Figure 1:**
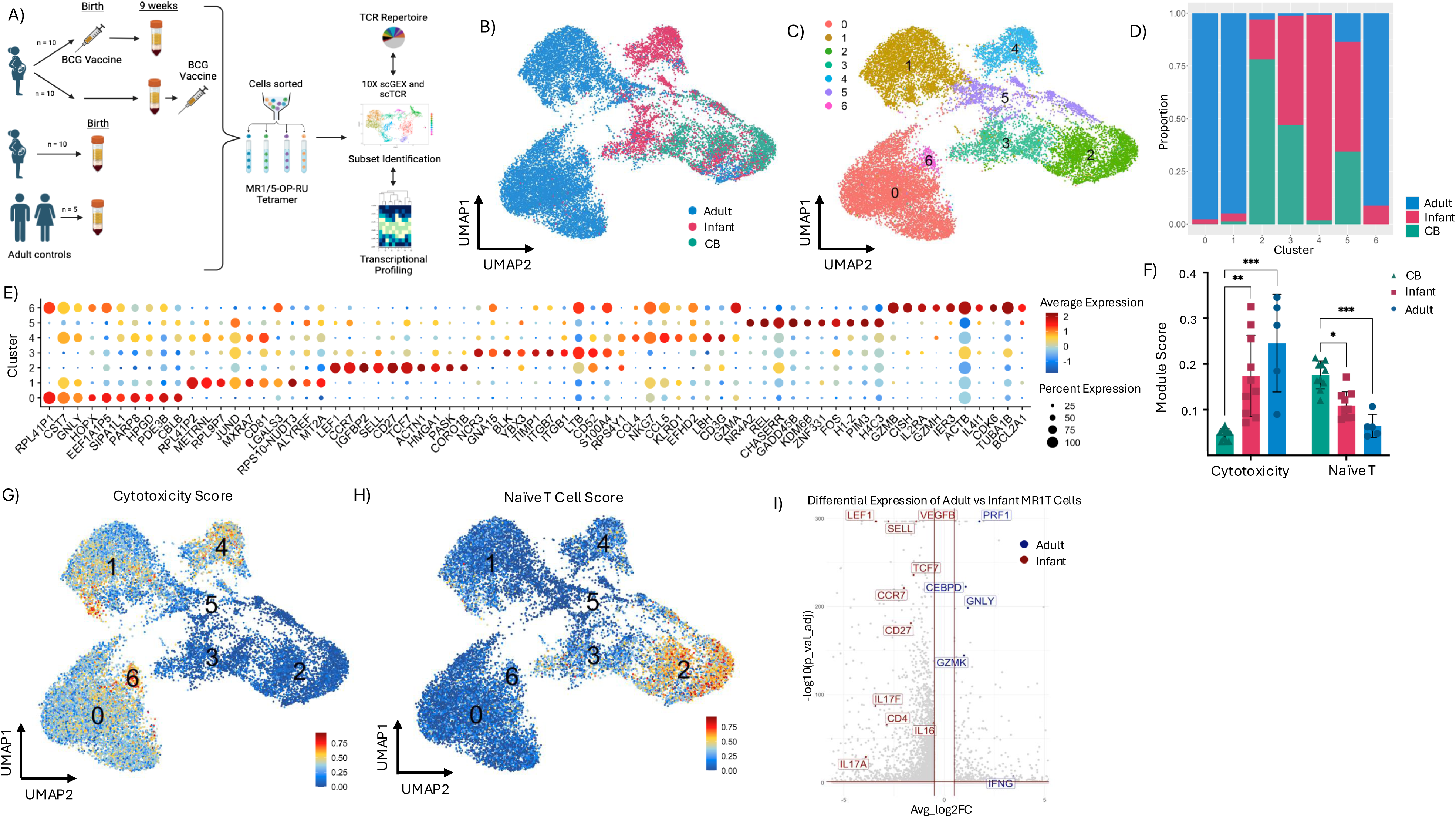
MR1T cells mature rapidly. A) Experimental workflow. scGEX = single-cell gene expression; scTCR = single-cell TCR. 5- OP-RU = 5-(2-oxopropylideneamino)-6-d-ribitylaminouracil. B) UMAP reduction of 5-OP-RU/MR1 sorted MR1T cells from cord blood, unvaccinated infant and adult donors. C) Unbiased Leiden clustering of MR1T cells. D) Proportions of each cluster composed of cord blood, infant or adult derived MR1T cells. E) Dotplot of top 10 genes upregulated in each cluster. Color represents average expression, and size represents percent of MR1T cells expression that gene. F) Cytotoxicity (*PRF1, GNLY, NKG7, GZMA, GZMB, GZMH, GZMK* and *GZMM*) and Naïve T (*CTSG, CA6, GSTT1, LEF1, RGS10* and *TMIGD2*) gene module scores for each donor segregated based on age of donor. Statistics done with Kruskal-Wallis. *p-value < 0.05; **p- value < 0.01; ***p-value < 0.001; ****p-value < 0.0001. Cytotoxicity CB vs infant p-value = 0.0014, cytotoxicity CB vs adult p-value = 0.0001, naïve CB vs infant p-value = 0.0187, and naïve CB vs adult = 0.0004. G) Cytotoxicity gene module overlaid on UMAP reduction with numbers representing Leiden clustering. H) Naïve T cell gene module score overlaid on UMAP reduction. I) Volcano plot of differentially expressed genes between infant (left; red) and adult (right; blue) donors. Red horizontal line shows the Bonferroni-adjusted p-value < 0.05 and red vertical lines represents -0.5 and 0.5 log fold change.

**Table 1.** Donor Characteristics.

|  | Cord Blood | Unvaccinated | BCG<br>Vaccinated | Adults | P-value<br>(Vaccinated vs<br>Unvaccinated) |
| --- | --- | --- | --- | --- | --- |
| Sex | 30% Male, 70%<br>Female | 60% Male, 40%<br>Female | 50% Male, 50%<br>Female | 80% Male, 20%<br>Female | 1.00 |
| Age | N/A | 63.2 days<br>(range 57-74) | 64.2 days<br>(range 57-71) | 29 years (range<br>24-32) | 0.668 |
| Gestational<br>Age (weeks) | Not recorded | Mean 39.5<br>(range 38-41) | Mean 38.4<br>(range 37-40) | N/A | 0.035 |
| Birth Weight<br>(kg) | Not recorded | 3.226 (range 2.9<br>– 3.98) | 3.048 (range<br>2.68 – 3.82) | N/A | 0.33 |

To examine the maturation of MR1T cells after birth, irrespective of BCG vaccination, cells from CB donors, unvaccinated infant donors and adult donors were clustered and UMAP reduction is shown in Figure 1B. Unbiased Leiden clustering yielded 7 distinct clusters (Figure 1C) with clusters 0, 1 and 6 enriched for adult MR1T cells, cluster 2 enriched for CB MR1T cells and cluster 4 enriched for infant MR1T cells (Figure 1D, Supplemental Figure 2A). The top 10 most upregulated genes in each cluster are shown in Figure 1E with cytotoxic genes (*CST7, GNLY, CCL4, CCL5, GZMA, NKG7, GZMB, GZMH*) upregulated in clusters 0, 4 and 6 and naïve and stem-like genes (*LEF1, CCR7, SELL, CD27, TCF7*) upregulated in cluster 2. This is also reflected in differences in cytotoxicity (*PRF1, GNLY, NKG7, GZMA, GZMB, GZMH, GZMK, GZMM*) and naïve T cell (*CTSH, CA6, GSTT1, LEF1, RGS10, TMIGD2*) UCell module scores,^34^ with significantly higher cytotoxicity scores in infant and adult donors and significantly upregulated naïve T cell scores in CB donors (Figure 1F). While adult versus infant cytotoxicity scores were not statistically significantly different, infant MR1T cells did trend towards a lower cytotoxicity score, and higher naïve T cell module score as compared to adults. Higher cytotoxicity module scores were seen in adult enriched clusters 0, 1, and 6 and in infant enriched cluster 4 (Figure 1G), whereas the naïve T cell module score was highest in CB enriched cluster 2.

Oligo-conjugated antibody staining for CD4, CD8, CD45RA, CCR7 and TRAV1-2 is shown in Supplemental Figure 2B. This corroborates the gene expression findings, showing increased expression of CD4 and CCR7 in cluster 2 (enriched for CB-derived MR1T cells) and upregulation of CD8, and TRAV1-2 in clusters enriched for adult and infant-derived MR1T cells. CD4 and CD8 expression on TRAV1-2(+) versus TRAV1-2(-) MR1T cells demonstrates increased CD4 expression on TRAV1-2(-) MR1T cells across the life span, although CD4 expression is most common amongst CB TRAV1-2(-) MR1T cells (Supplemental Figure 2C). Among TRAV1-2(+) MR1T cells, CB and infant-derived MR1T cells demonstrated higher CD4 expression, while adult MR1T cells are comprised of predominately CD8 expressing MR1T cells.

Highlighting the rapid maturation of MR1T cells after birth, MR1T cells from infants at 9-weeks of age had significant upregulation of inflammatory and cytotoxic genes (*PRF1, NKG7, GNLY, CCL5, CCL4, KLRB1, KLRD1, GZMA, GZMB, GZMK, GZMH, IL32, KLRG1, FASLG, IL26, IL18R1, CXCR4, KLRC1, CXCR3, TNF, IFNG, CCR5, CCL20, IL12RB1*) compared to CB MR1T cells (Supplemental Figure 3A, Supplementary Table 1). Moreover, infant MR1T cells demonstrated upregulation of tissue homing genes (*CEBPD, CXCR6, CXCR3*), canonical TCR genes (*TRAV1-2),* along with critical transcription factors associated with effector status (*EOMES, TBX21, ZBTB16*). CB cells meanwhile had significant upregulation of naïve, repair and stem-like genes (*LEF1, TCF7, CCR7, CD27, SELL, VEGFB, IL16, CD4*) compared to infants (Supplemental Figure 3B, C). The differentially expressed genes in infant MR1T cells were similar to those upregulated in adult MR1T cells (Supplemental Figure 3A-C, Supplementary Table 2) with many of the same genes upregulated and down regulated compared to CB donors. Although there are many shared upregulated genes in infant and adult MR1T cells, adult MR1T cells did have further upregulation of unstimulated cytotoxic gene expression (*PRF1, GNLY, GZMK, IFNG*), along with the tissue residency gene *CEBPD* in adult MR1T cells (Figure 1I, Supplementary Table 3).

### MR1T Cells Have Diverse TCR Usage at Birth, that becomes more focused on TRAV1-2(+) with aging

To explore how TCR usage changes over early life, single-cell TCR sequencing (scTCR-seq) was performed at the same time as scRNA-seq. At birth, CB-derived MR1T cells had a diverse TCR repertoire with the majority of cells utilizing TRAV1-2(-) α-variable chains (Figure 2 A, B). Even by 9 weeks of age we observed the rapid expansion of the TRAV1-2(+) subset. Adult MR1T cells demonstrate additional skewing towards TRAV1-2 usage based on TCR-sequencing (Figure 2 A, B, C). Figure 2D shows further analysis of TCR usage using scTCR-seq. CB-derived MR1T cells were also much more diverse in their TRAJ, CDR3α and TRBV usage compared to infant-derived MR1T cells. In addition to increasing TRAV1-2 usage we also observe increased usage of the canonical TRAJ33 with age. Using CDR3α sequences to denote unique α chains, one canonical CD3α increased modestly (CAVMDSNYQLIW) from birth to infancy and adulthood, while another canonical CD3α (CAVRDSNYQLIW) expanded dramatically in infants and adults as compared to CB. Similarly, canonical TRBV (TRBV6-4, TRBV6-1, TRBV20-1) usage increased from representing approximately a quarter of CB MR1T cells to more than half in infant and adult MR1T cells. Increased TCR diversity with age is also demonstrated by comparison of CDR3α and CDR3β among CB, infant, and adult MR1T cell TCR’s (Supplemental Figure 4). Specifically, the CDR3α length is much more variable for CB-derived MR1T cells, compared to infant or adult MR1T cells (Supplemental Figure 4A). Adult MR1T cells were nearly uniform in length (12 amino acids). CDR3β length was more variable in all age groups, reflecting the diversity of β-chains for MR1T cells. The major factor influencing CDR3α length was TRAV1-2 usage, whereas TRAV1-2 usage did not influence CDR3β length when examining all cohorts combined (Supplemental Figure 4B). Finally, due largely to β chain diversity, even among TRAV1-2(+) MR1T cells, the majority of TCRs were private to individual donors (Supplemental Figure 4C).

**Figure 2:**
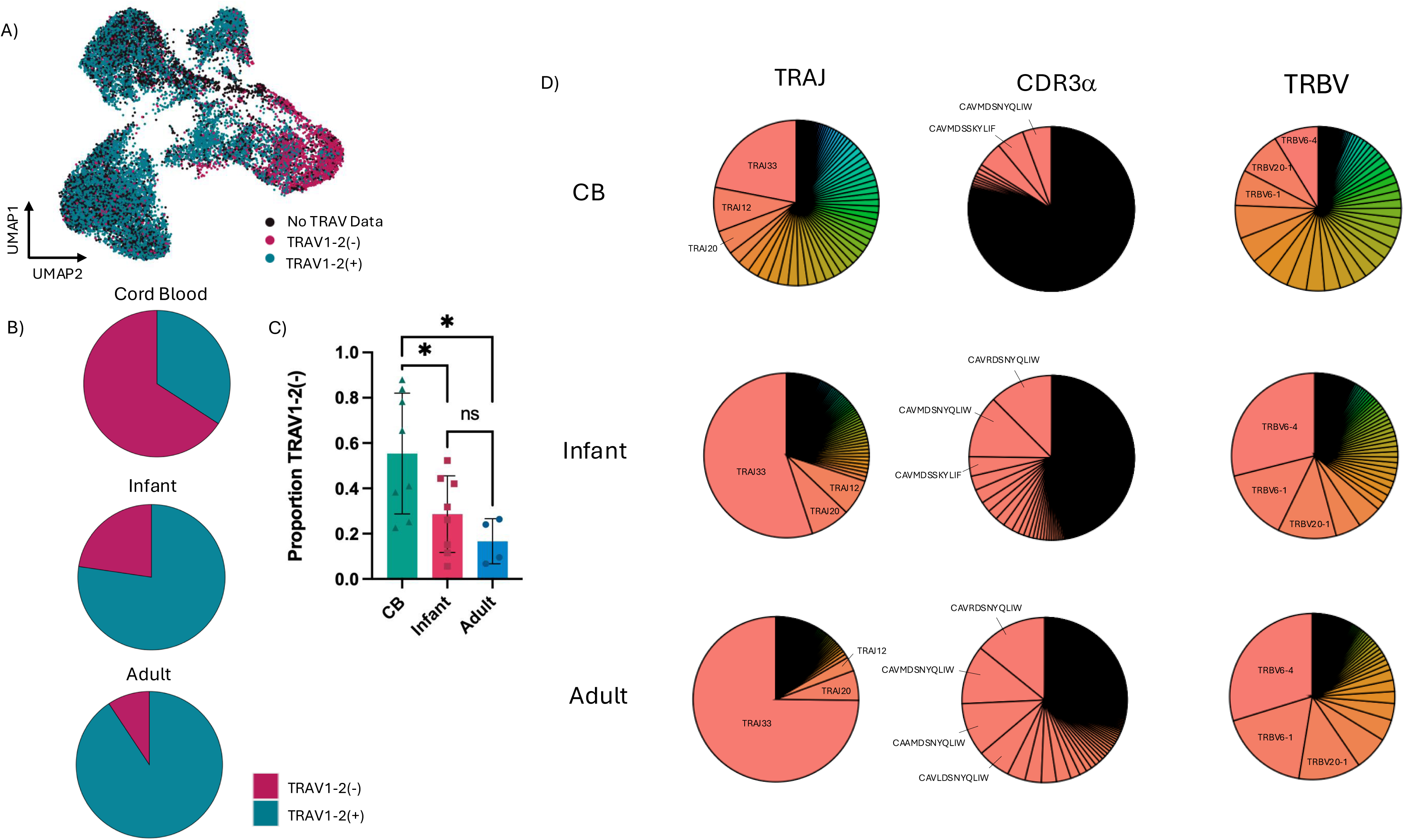
Diverse TCR usage at birth, becomes adult-like by 9-weeks of age. A) UMAP reduction showing TRAV1-2(+), TRAV1-2(-) or cells with no TRAV data based on TCR-sequencing. B) Proportion of cord blood, infant or adult derived TRAV1-2(+) or TRAV1-2(-) MR1T cells (excluding cells which do not have TRAV data). C) Individual proportion of each donor’s cells that are TRAV1-2(-) for all donors with at least 100 cells with TCR sequences. *p-value < 0.05; one-way ANOVA and Tukey’s HSD. CB vs infant p-value = 0.0483 and CB vs adult p-value = 0.0188. D) TRAV, TRAJ, CDR3α and TRBV usage for cord blood, infant and adult MR1T cells. Top values are labelled. TRAV1-2(+) cells that also expressed a second TRAV are combined under the label TRAV1-2,TRAV(X).

### TRAV1-2(+) MR1T Cells are Less Naïve, Even at Birth

To examine if there were differences in the maturation of TRAV1-2(+) compared to TRAV1-2(-) MR1T cells, common naïve and effector genes were examined for each donor stratified based on TRAV1-2 usage (Figure 3A-D). In CB donors, both TRAV1-2(+) and TRAV1-2(-) MR1T cells showed low effector gene modules scores and were not significantly different, however, TRAV1-2(+) cells had significantly lower naïve T cell module scores. This difference was maintained in infant MR1T cells with significant reductions in naïve T cell module scores in TRAV1-2(+) cells. There was also an increased cytotoxicity gene module score for TRAV1-2(+) cells in infants and adults, but this was not significant. For adult MR1T cells there were no longer significant differences in naïve T cell module score, with all cells having a relatively lower module score compared to earlier in life.

**Figure 3:**
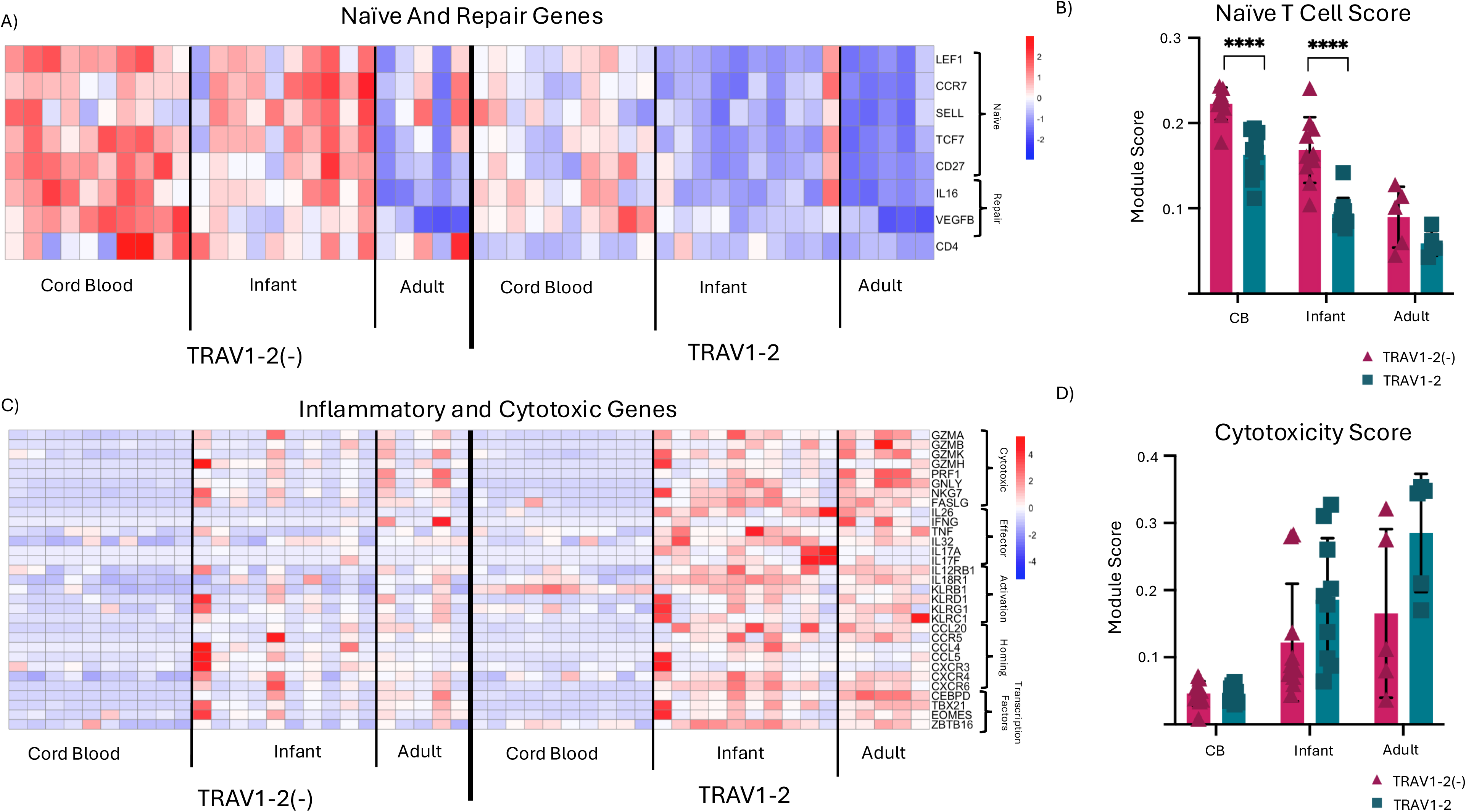
Even in Cord Blood, TRAV1-2(-) are more naïve. A) Average expression of naïve and repair genes for each individual donor. B) Naïve T module score (*CTSG, CA6, GSTT1, LEF1, RGS10* and *TMIGD2*) for each donor split by TRAV1-2(+) compared to TRAV1-2(-) MR1T cells. **p-value < 0.01; ***p-value < 0.001; ****p-value < 0.0001; student t-test. CB TRAV1-2(+) vs TRAV1-2(-) p-value = <0.0001, infant TRAV1-2(+) vs TRAV1-2(-) p-value = <0.0001. C) Average expression cytotoxic and pro-inflammatory genes for each individual donor. D) Cytotoxicity module score (*PRF1, GNLY, NKG7, GZMA, GZMB, GZMH, GZMK,* and *GZMM*) for each individual donor. **p-value < 0.01; student t-test.

### BCG Vaccination Leads to Upregulation of Interferon-Stimulated Genes in MR1T Cells

Previous work by Gela *et al*,^32^ looked for changes in DURT frequency between BCG vaccinated, compared to unvaccinated 9-week-old infants and found no differences in frequencies of MR1T cells. This study, however, did not investigate if there were changes in MR1T cell function or clonal expansion of specific clonotypes that were modulated by newborn BCG vaccination. There are very significant changes to MR1T cells that occur in early life which may mask more subtle shifts caused by BGC vaccination. To further explore this, we directly compared MR1T cells from infants who had received BCG vaccine at birth, compared to infants who had delayed BCG vaccination until after blood was collected at 9-weeks of age (Figure 1A). These infant MR1T cells were re-clustered and UMAP reduction is shown in Figure 4A. Differential gene expression comparing vaccinated to unvaccinated infants showed significant upregulation of interferon-stimulated genes (ISG) in vaccinated infants (Figure 4B, Supplemental Figure 5A-D, Supplementary Table 4). This is reflected in the top GO pathways upregulated in vaccinated infants associated with response to interferons and defense against viruses (Figure 4C). Unbiased Leiden clustering of infant MR1T cells resulted in 9 distinct clusters (Figure 4D). Most clusters have similar distribution between vaccinated and unvaccinated infants, but cluster 6 was enriched for MR1T cells from vaccinated infants (Figure 4E). This cluster also demonstrated the highest interferon response (*IFI6, IFI27, MX1, ISG15, STAT1, LOC114672189, MX2, IFIT3*) gene module score (Figure 4F). Differential gene expression of cluster 6 compared to other clusters showed significant upregulation of ISG (*IFI6, IFI44L, MX1, LY6E, MX2, TIM22, IFI35, OAS3, OASL, HERC5, ISG15, OAS1, IFIT1, PLSCR1, IFIT3*) but also significant upregulation of cytotoxic genes (*NKG7, GZMK, GZMA, GZMH, GZMB, KLRG1, KLRD1, CCR5, CCL4, CCL5*), critical transcription factors associated with effector status (*EOMES, TBX21*) and tissue homing genes (*CEBPD, LGALS3BP*) (Figure 4G, H, Supplemental Figure 6A, B, Supplementary Table 5).

**Figure 4:**
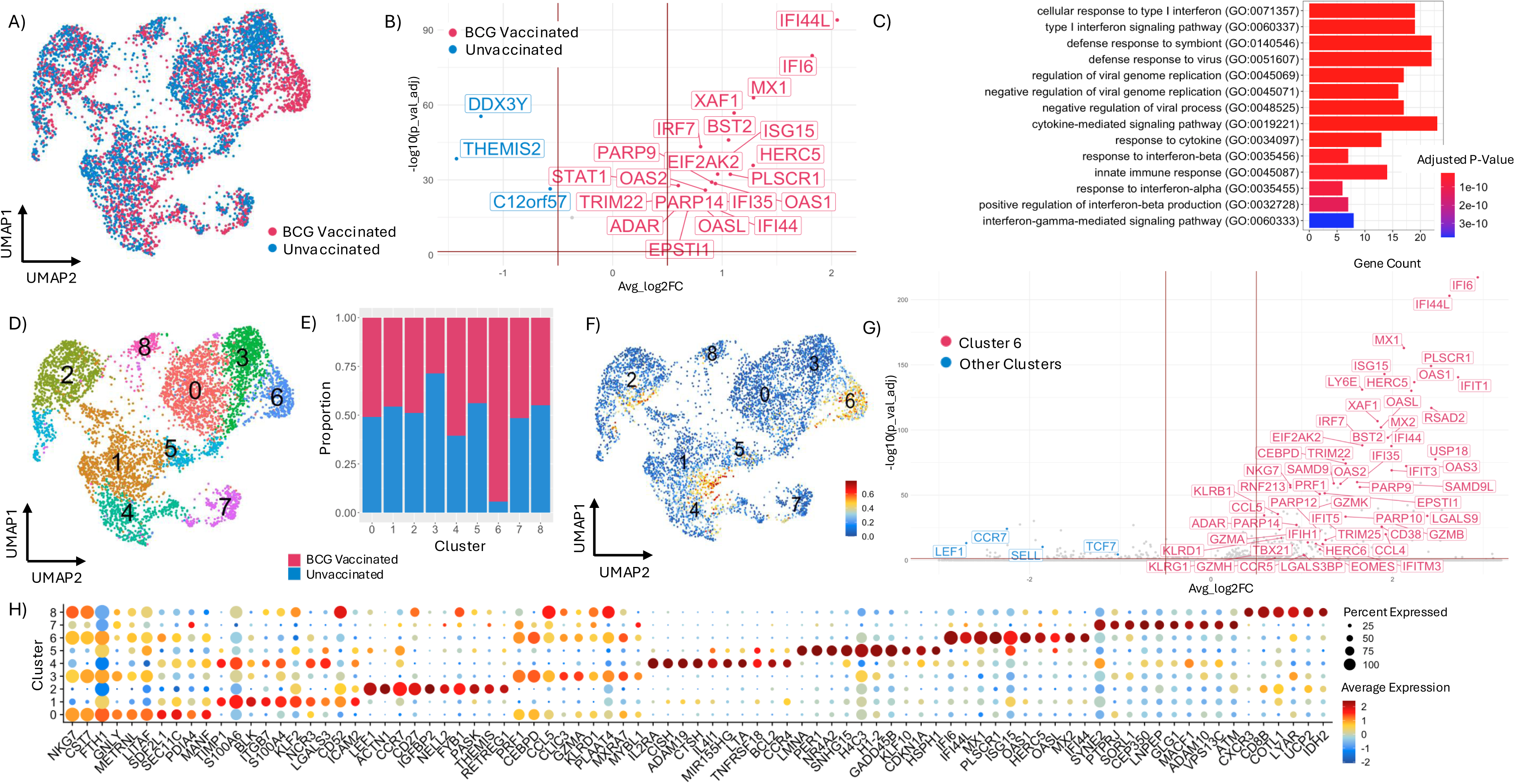
Enrichment of interferon stimulated genes in MR1T cells following BCG vaccination. A) UMAP reduction of BCG vaccinated (pink) and unvaccinated (blue) infant MR1T cells. B) Volcano plot of differentially expressed genes between BCG vaccinated (pink; right) and unvaccinated (blue; left). Red horizontal line showing Bonferroni-adjusted p-value < 0.05 and red vertical lines representing -0.5 and 0.5 logfold change. C) Top upregulated GO pathways of differentially expressed genes in BCG vaccinated infant MR1T cells compared to unvaccinated infant MR1T cells. D) Unbiased Leiden clustering of infant MR1T cells. E) Proportion of each cluster composed of BCG vaccinated vs unvaccinated infant MR1T cells. F) Interferon response (*IFI6, IFI27, MX1, ISG15, STAT1, LOC114672189, MX2, IFIT3*) gene module score shown over UMAP reduction with number representing Leiden clustering. G) Volcano plot of Leiden cluster 6 (pink; right) compared to other clusters (blue; left). H) Dotplot of top 10 genes upregulated in each cluster. Color representing average expression and size representing percent of MR1T cells expression that gene.

Cell surface protein expression was similar between vaccinated and unvaccinated infants (Supplemental Figure 6B, C). The majority of infant MR1T cells expressed canonical cell surface protein markers including TRAV1-2, CD8, CD26 and CD161. There remained a smaller subset of non-canonical CB-like cells that did not stain for TRAV1-2 and were predominantly CD4 expressing. This is reflected in Supplemental Figure 6D, where cells were gated based on cell surface protein expression for CD4 and/or CD8. For both vaccinated and unvaccinated infants, TRAV1-2(+) cells were more likely to be CD8 or DN compared to TRAV1-2(-) cells, which had a higher frequency of CD4 expressing MR1T cells. Overall distribution of CD4/CD8 status was not significantly changed by BCG vaccination (Supplemental Figure 6E).

### BCG Vaccination Does Not Change MR1T TRAV, TRAJ, or TRBV Usage

We next investigated whether BCG vaccination modulates MR1T TCR usage. There were no major shifts in utilization of common TRAV, TRAJ or TRBV between BCG vaccinated and unvaccinated infants (Supplemental Figure 7). Given the predominant expression of the canonical TRAV1-2 α chain by 9-weeks of age we were not surprised to also observe little effect of BCG on α chain diversity (Fig 5A, B, Supplemental Figure 7). By contrast, we observed that the majority of CDR3β-chains were unique to an individual and thus very different between BCG vaccinated and unvaccinated infants (Figure 5C, D). Given the privacy of CDR3β-chains, it’s not useful to directly compare frequencies of individual CDR3β-chains, but what we did observe is that there were fewer unique expanded CDR3β pairings in BCG vaccinated infants, defined as > 3 CDR3β-chains (Figure 5D). This suggests the expansion of BCG-selective TCRs.

**Figure 5:**
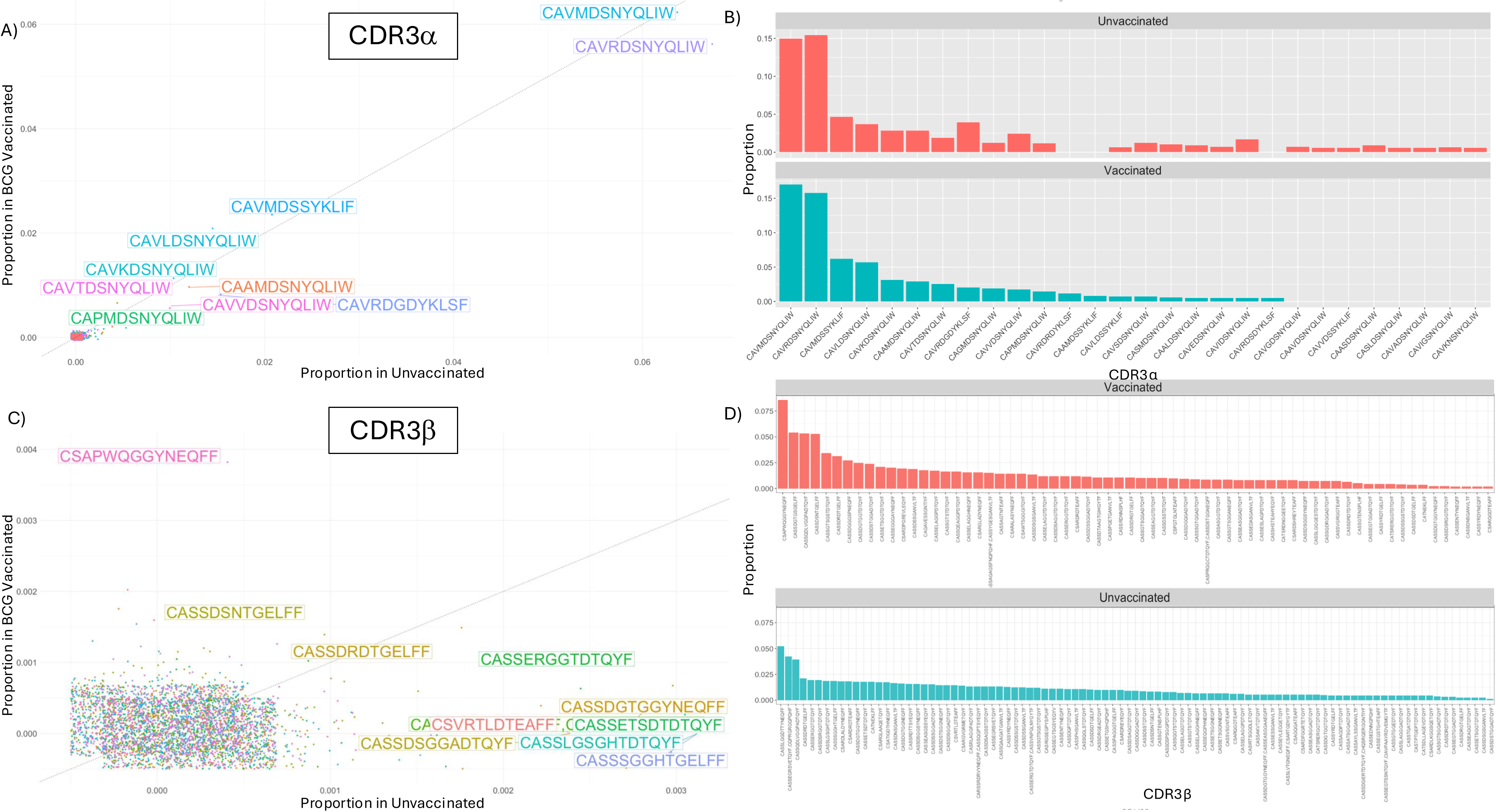
Infant MR1T cells have predominantly public CDR3α and predominantly private CDR3β. A) Frequency of CDR3α in BCG vaccinated (y-axis) compared to unvaccinated (x-axis) infants. Highest frequency CDR3α are labeled. Proportions are based on total cells. B) Bar plot of most frequent CDR3α with proportions based on only cells with available CDR3α data. C) Frequency of CDR3β in BCG vaccinated (y-axis) compared to unvaccinated (x-axis) infants. Highest frequency CDR3β are labeled. Proportions are based on total cells. Points at 0 are jittered to display density. D) Bar plot of expanded CDR3β sequences (> 3 identical sequences) with proportions based on only cells with available CDR3β data.

### Expanded Clonotypes in Infant-derived MR1T Cells are Less Naïve and More Cytotoxic

We next examined expanded clonotypes (≥ 3 identical CDR3 regions) in the infant samples to determine if these expanded clonotypes were distinct from unexpanded clonotypes. Expanded clonotypes were more frequently found in clusters with higher cytotoxicity score (Figure 6A, B) and had a higher cytotoxicity score compared to low frequency clonotypes (< 3 identical CDR3 regions found) (Figure 6C). Compared to expanded clonotypes, for which top upregulated genes include cytotoxic and effector genes (*PRF1, GZMA, IL18RAP*), low frequency clonotypes upregulated naïve and stem-like genes including *CCR7* and *LEF1* (Figure 6D).

**Figure 6:**
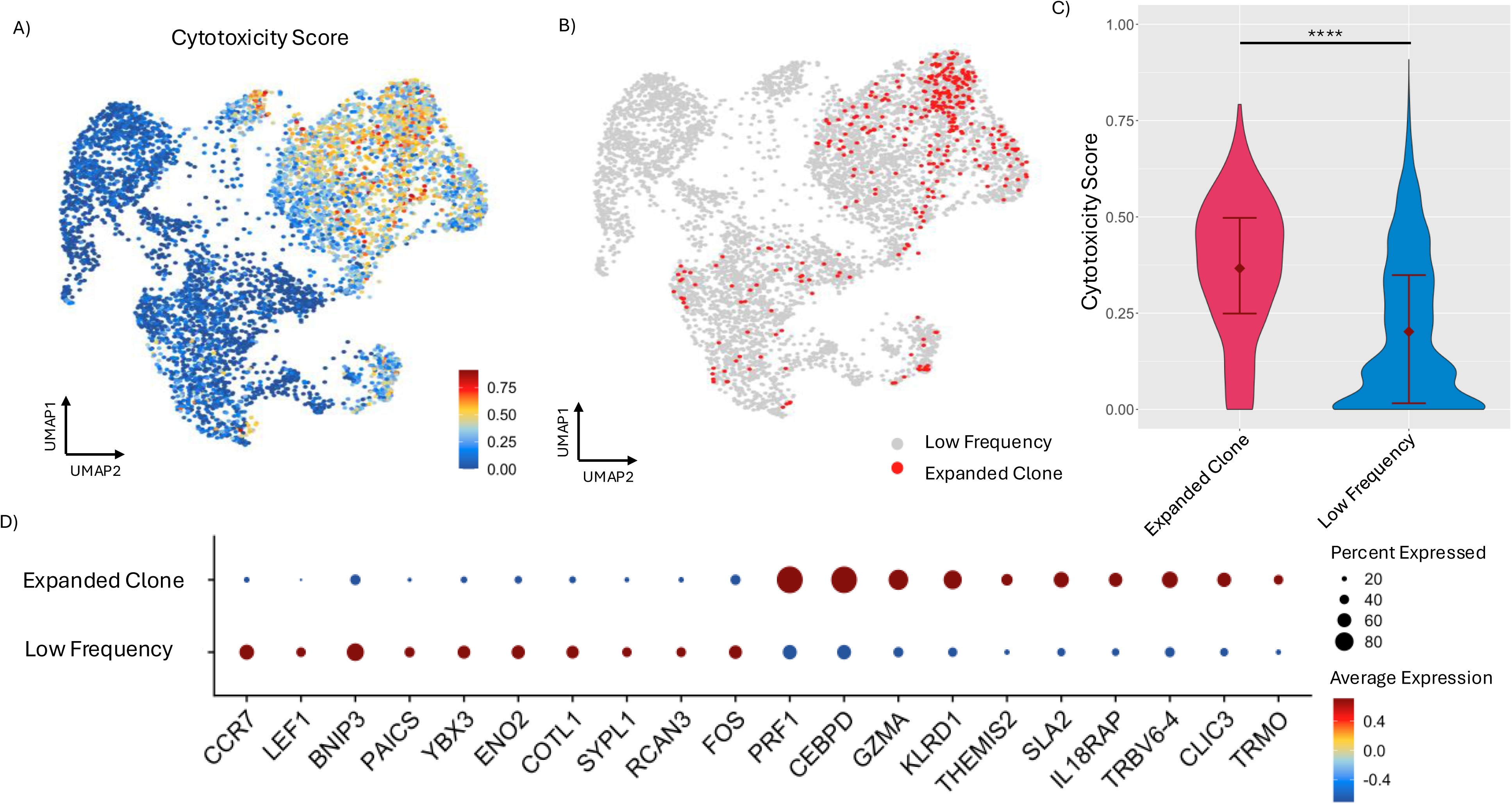
Expanded clonotypes upregulate cytotoxic genes and downregulate naïve genes. A) Cytotoxicity module score (*PRF1, GNLY, NKG7, GZMA, GZMK,* and *GZMM*) overlaid on UMAP dimensionality reduction of infant MR1T cells. B) UMAP reduction showing expanded clone (≥ 3 identical CDR3 regions). C) Cytotoxicity module score for expanded clones compared to low frequency clones. ****p-value < 0.0001; Welch’s t-test. D) Dotplot of top 10 genes upregulated in expanded clonotypes comparted to low frequency cells. Color representing average expression and size representing percent of MR1T cells expression that gene.

### BCG Vaccination Leads to Selective Expansion of a TCR Cluster

To evaluate if BCG vaccination induces selective expansion of MR1T cells with shared or common TCR features, we compared MR1T cells between the BCG vaccinated versus BCG unvaccinated infants utilizing the TCR clustering algorithm TCRdist3 to look for enriched TCR clusters in vaccinated infants. TCRdist3 was used to cluster only CDR3β chains, rather than the paired CDR3α/β. This was done because of the semi-invariant α-chain in MR1T cells, to prevent over-representation of the more diverse TRAV1-2(-) MR1T cells that become rare after birth. In total there were 19 TCR clusters (Figure 7A). When comparing the frequencies of cells within each TCR cluster between BCG-vaccinated and unvaccinated infants, only Cluster 13 showed a significant difference, with a marked enrichment in BCG-vaccinated infants (p = 0.0012). This remained significant when the proportion of each infant in cluster 13 was analyzed (Figure 7B). Differential gene expression of TCR cluster 13 compared to other TCR clusters showed 31 significantly upregulated genes in cluster 13 (Figure 7C, Supplementary Table 6). These include effector and cytotoxic genes (*GZMM, NKG7, GNLY, CCL5, CD8A, CD8B, CD96, TNFAIP3*), tissue residency and homing (*KLRB1, CXCR6, SELPGL, CEBPD*), along with TCR signaling and T cell homeostasis (*IL7R, FYN, CD6, RORA, MT2A, TNIK*). Cluster 13 also downregulated 11 genes including the naïve T cell marker: *CCR7*.

**Figure 7:**
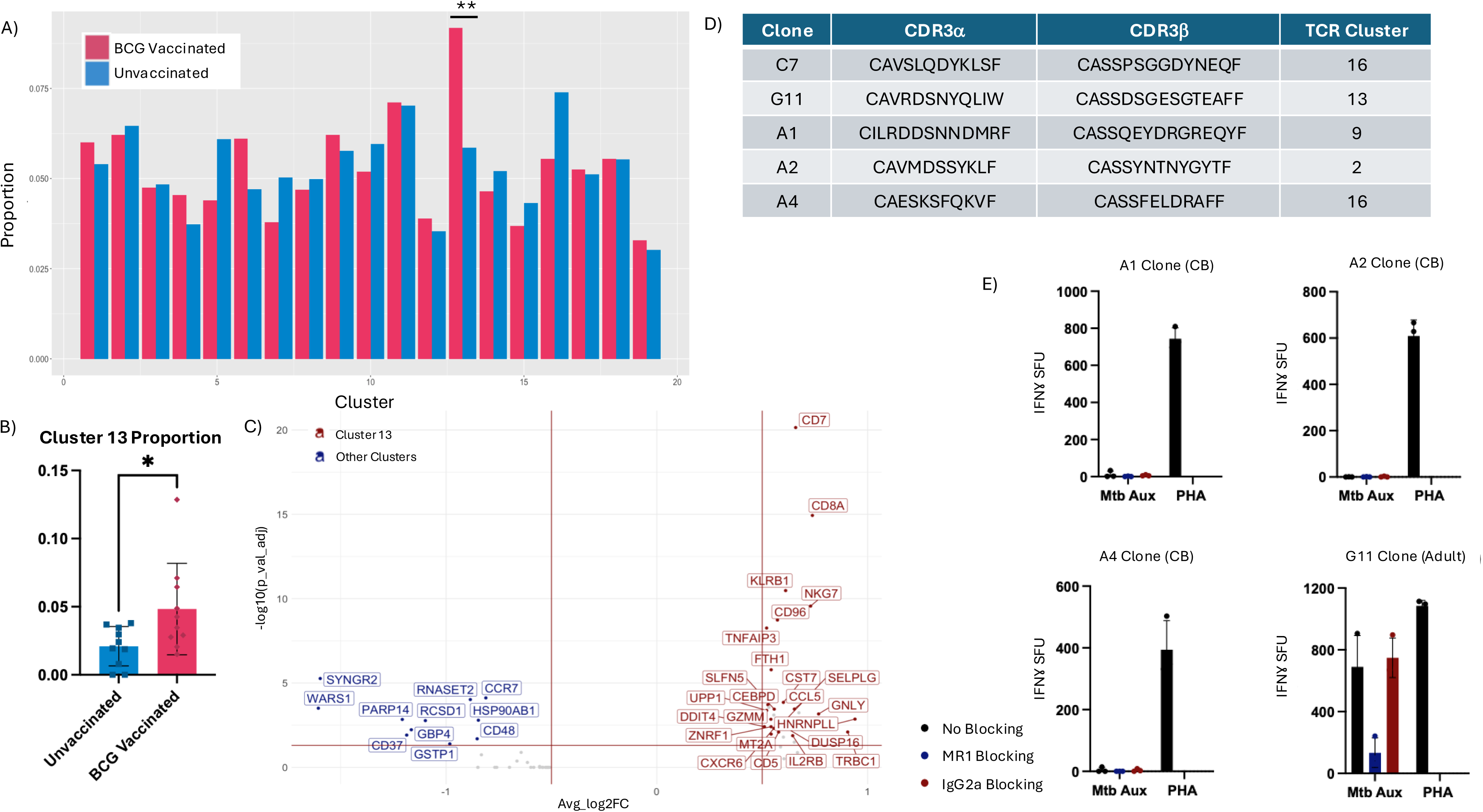
BCG Vaccination Leads to Expansion of Specific TCR Cluster. A) Proportion of total BCG-vaccinated (pink) or unvaccinated (light blue) infant cells that make up each TCRdist3 cluster. P-values calculated using Fischer exact test. ** < 0.01, non-significant values denoted with no markings. B) Proportion of each individual donor’s cells that fall into cluster 13 of the TCRdist3 clusters. p-value = 0.0195; Wilcoxon test. C) Differentially expressed genes in TCRdist3 cluster 3 compared to other TCRdist3 clusters. Red horizontal line showing Bonferroni-adjusted p-value < 0.05 and red vertical lines representing -0.5 and 0.5 log fold change. D) CDR3 regions for C7 and G11 clones previous described.^31,44^ E) IFNγ ELISPOT of CB MR1T cell clones (A1, A2 and A4) and adult MR1T cell clone (G11) response to Mycobacteria and Mycobacterial antigens. Responses without blocking, anti-MR1 blocking antibody or isotype control blocking antibody are represented. Previously published in Kain *et al.*^44^ SFU = spot forming units. PHA = phytohaemagglutin P. PL1 = photolumazine I. DZ = deazalumazine.

To help determine the mycobacterial-reactivity of TCR cluster 13, enriched in BCG vaccinated infants, we utilized TCR sequences previously derived from two adult MR1T cell clones, G11 and C7 (recognize Mtb), and three cord blood derived MR1T cell clones: A1, A2 and A4 (do not recognize Mtb).^31,35^ The TCR sequences of these clones are shown in Figure 7C, and these sequences were spiked into the TCRdist3 analysis, finding that G11 falls within cluster 13, while the other clones fall into other clusters (Figure 7C). G11, A1, A2 and A4 clones were analyzed for response to different mycobacterial antigens in an IFNγ ELIspot as previously published (Figure 7D).^35^ The A1, A2 and A4 clones do not respond to Mtb auxotroph, whereas G11 is Mtb responsive, suggesting expansion of a mycobacterial-reactive cluster with BCG vaccination.

## Discussion

Perinatal BCG vaccination has been shown to reduce the risk of severe and disseminated TB in early life but does not provide consistent benefit against TB prevention in adults.^6^ The reasons for these differences remain unknown. Studying the effect of BCG in early life and its role on the developing immune system could provide critical insights into BCG-mediated protection from TB. BCG has been shown to induce a robust CD4 T cell response in NHP^36^ and human studies,^37,38^ and these have long been the focus on TB vaccine development. Other branches of the immune system also recognize BCG but have been understudied. This includes groups of DURTs such as MR1T cells, which are able to robustly respond to mycobacteria and produce important cytokines to defend from Mtb, including IFNγ^39^ and TNF,^40^ as well as cytotoxic molecules that are increasingly recognized to be important in protection.^29,41,42^ Here we show that BCG leads to both an interferon-stimulated gene signature in MR1T cells and induces selective expansion of a mycobacterial-reactive TCR cluster.^33,43^ This work demonstrates that BCG-vaccination shapes the selective expansion of MR1T cells and suggests they could potentially be harnessed for future vaccines.

It has previously been shown that MR1T cells acquire effector functions, including the ability to produce IFNγ, in the developing thymus.^44^ Despite this, MR1T cells at birth have very distinct gene expression profiles from adult MR1T cells with a naïve phenotype and an extremely diverse TCR repertoire.^45^ There is a rapid shift, however, over the first few months of life, to an adult-like phenotype and increased functional ability of MR1T cells.^16^ Our work here confirms these results but adds additional context to infant MR1T cells, finding that although they appear similar to adult MR1T cells, they undergo further maturation later in life, as evidence with downregulation of naïve genes and upregulation of cytotoxicity genes. Notably, this later maturation is particularly seen in TRAV1-2(-) MR1T cells. Prior studies have shown that TRAV1-2(-) MR1T cells are less responsive to common childhood pathogens such as *S. aureus, S. pneumoniae,* and *M. tuberculosis*,^45^ and in some cases may recognize non-riboflavin bacteria such as *S. pyogenes*.^15^ These findings raise the possibility that the slower maturation trajectory of TRAV1-2(-) MR1T cells reflects differential microbial exposures that selectively drive their expansion and functional programming.

Previous work in adults has shown that BCG revaccination leads to a transient increase in peripheral MR1T cell frequencies, with MR1T cells representing the predominant BCG-reactive IFNγ-producing CD8 T cell.^46^ This response was however largely TCR-independent and instead cytokine mediated. In contrast, BCG vaccination at birth did not lead to a similar expansion of peripheral MR1T cells.^32^ Consistent with this, BCG vaccination in mice also does not lead to peripheral MR1T cell expansions but did when combined with a TLR2/6 agonist.^47^ Importantly, MR1T cells appear functionally relevant in the murine response to BCG, as MR1⁻/⁻ mice demonstrate impaired control of experimental *M. bovis* infection.^48^ These observations raise the possibility that, despite the absence of frequency changes in infants, BCG vaccination may still induce functional alterations in MR1T cells that contribute to protection.

Supporting this, our data demonstrate a significant upregulation of ISGs in infant MR1T cells at 9 weeks post-BCG vaccination. MR1T cells are known to respond to type I interferons (IFN-α/β) in a TCR-independent manner, enhancing their capacity to produce IFN-γ and cytotoxic effector molecules.^49^ Indeed, López-Rodríguez *et al.* also showed that MR1T cell-mediated control of *Klebsiella pneumoniae* pulmonary infection in mice depended on type I interferon signaling.^50^ In our study, the lack of overlap between expanded MR1T clonotypes and ISG-enriched transcriptional clusters further supports a TCR-independent, cytokine-driven mechanism. In NHP models, IV BCG induced an early strong ISG signature, and this was associated with protection from protection from Mtb challenge.^51^ Together, these findings suggest that BCG vaccination at birth induces type I interferon responses that may exert prolonged cytokine-mediated effects on MR1T cells, potentially priming them for enhanced responses to Mtb or other neonatal pathogens. Given the live attenuated nature of BCG, and its capacity for prolonged persistence in infants, it remains unclear whether the observed gene expression changes reflect ongoing antigenic stimulation from persistent live BCG or downstream effects of early BCG-driven innate immune activation.

Attempts to harness MR1T cells through vaccination with 5-OP-RU have been largely unsuccessful in both mouse and non-human primate models. In mice, vaccination with 5-OP-RU and using cytosine-phosphorothioate-guanine (CpG) as an adjuvant led to delays in T cell priming and failed to provide protection against subsequent Mtb challenge.^52^ Similarly, in non-human primate models, 5-OP-RU vaccination with CpG led to MR1T cells upregulating PD-1 and losing their functional ability to produce many important cytokines.^52^ MR1T cells have, however, been shown to have more antigen selectivity that previously thought, with several studies showing selective antigen responses in MR1T cells.^30,31^ Therefore, despite these disappointing early results, it’s possible that 5-OP-RU vaccination is not the best strategy to harness these cells. This antigen may be too broad, activating MR1T cells in a non-specific manner and leading to exhaustion of these cells, rather than protection. Furthermore, recent work has shown the plasticity of MR1T cells, wherein different cytokines at the time of activation can shift IL-17 producing MR1T cells to become IFNγ producers.^53^ This flexibility suggests that MR1T cells are not irreversibly committed to a single functional program but instead retain the capacity to adapt to their surrounding environment. Such plasticity raises the alternative hypothesis that the cytokine milieu induced by different adjuvants could critically shape the functional trajectory of MR1T cells following vaccination. In this context, it is also possible that longer or repeated stimulation with live attenuated BCG, which more closely mimics natural infection, may be necessary to drive durable programming of MR1T cells toward protective effector functions.

Our findings support this hypothesis by demonstrating that BCG vaccination does not broadly expand the MR1T cell repertoire, but instead selectively expands specific TCRs. The lack of major changes in the TCR repertoire with vaccination is not unexpected given the large expansions in early life driven by the microbiota.^54^ The noticeable reduction in the diversity of expanded MR1T CDR3β regions in BCG vaccinated infants is significant as the CDR3β has been demonstrated to be very important in antigen discrimination.^10,55–57^ Further supporting the hypothesis that BCG leads to the selective expansion of specific mycobacterial-reactive-TCRs, there was expansion of a single TCR-cluster in BCG-vaccinated infants and this cluster upregulated many important effector genes. This includes critical tissue residency genes in this cluster (*CXCR6, SELP1G, CEBPD*), suggests that these expanded populations may be more able to take up tissue residency and provide protection at barrier sites. In particular, *CEBPD* expression has been shown to be critical for MR1T cells to exit blood vessels and enter the tissues,^58^ such as the lungs. Overall, this work suggests that BCG vaccination at birth leads to an expansion of a mycobacterial-reactive TCR cluster with upregulation of genes that may make these cells more adept at response to secondary exposures.

By comparing infants who had received BCG vaccination at birth, or who had BCG vaccination delayed, we are directly able to observe how MR1T cell ontogeny is influenced by a defined mycobacterial exposure. We found that BCG-vaccinated infant MR1T cells upregulated interferon stimulate genes, and through utilization of TCR clustering algorithms, showed that there are specific MR1T cell expansions in response to BCG-vaccination. This work has important implications for both understanding the development of MR1T cells in early life, but also their ability to be harness for novel vaccine development.

## Methods

### Study Participants

This study was conducted according to the principles expressed in the Declaration of Helsinki. Study participants, protocols, and consent forms were approved by the Institutional Review Board of the University of Cape Town (refs: 126/2006, 479/2009, 177/2011) for infant and CB donors, and at the University of Stellenbosch for adult donors (MRM study SUN HREC approval number 7 study title: N19/07/093; “MR1 Restricted memory T cells and TB”). All ethical regulations relevant to human research participants were followed. Umbilical CB was obtained after delivery and collected into citrate CPT tubes (BD) from the placenta of uncomplicated term pregnancies in Cape Town region, South Africa. Venous blood samples were collected into sodium heparin from 9-week-old infants and processed in a similar manner, as were adult samples all from participants in Cape Town region, South Africa. Blood was processed per manufacturer’s instructions within 24 hours of collection and then resulting CBMC was cryopreserved. Samples from the same cohort were previously used for another study.^32^

### Single-cell Sequencing of MR1T Cells and MR1T Cell Clones

The following reagents were obtained through the NIH Tetramer Core Facility: MR1/5-OP-RU tetramer and MR1/6- formylpterin (6-FP) tetramer.^59^ Identification of MAIT cells with MR1 tetramers was described previously.^11,16,35^ CMBC or PBMC was thawed in the presence of DNase, resuspended in 10% heat inactivated human serum with RPMI (Lonza) at a concentration of 2x10^7^/ml. CBMC/PBMC (2 x 10^6^ cells/well) was then stained with MR1/5-OP-RU tetramer at a dilution of 1:250 for 45 minutes at room temperature.^11,16,35^ Cells were then subsequently stained with an antibody cocktail of surface stains (listed in Supplemental Table 7) for 30 min at 4°C to allow non-CD3+ T cells or dead cells to be excluded. Cells were incubated in Human TruStain FcX (Biolegend^TM^) for 10 minutes at 4°C and then stained with TotalSeq-C hashtag antibodies and CITE-seq antibodies (Biolegend^TM^) listed in Supplemental Table 8 for 30 minutes at 4°C. Samples were then then sorted using a BD FACSAria^TM^ III Cell Sorter for MR1/5-OP-RU tetramer positive cells (representative gating strategy shown in Supplemental Figure 1). Sorted MR1T cells were then loaded in 10X Genomics^TM^ Chromium Next GEM Single Cell 5’ v2. Full protocol details of single-cell GEX, TCR and cell surface protein library preparation can be found from 10X genomics website (https://cdn.10xgenomics.com/image/upload/v1666737555/support-documents/CG000331_ChromiumNextGEMSingleCell5-v2_UserGuide_RevE.pdf). Finished libraries were then sent to Novogene Corporation Inc.^TM^ in Sacramento California for NovaSeq 6000 sequencing. MR1T cell clones were also sequenced using 10X Genomics^TM^ Chromium Next GEM Single Cell 5’ v2.

### Single-Cell RNA-seq Pre-processing

Raw sequence reads were processed using 10X Genomics Cell Ranger software (version 6.1.1). The resulting sequence data were aligned to the GRCh38 human genome. Cell demultiplexing used a combination of algorithms, including GMM-demux, demuxEM and BFF, implemented using the cellhashR package.^62–63^ Droplets identified as doublets (i.e. the collision of distinct sample barcodes) were removed from downstream analyses. We additionally performed doublet detection using DoubletFinder, and removed doublets from downstream analysis.^63^ Next, droplets were filtered based on UMI count (allowing 0-20,000/cell), and unique features (allowing 200-5000/cell). Additionally, we computed a per-cell saturation statistic for both RNA and ADT data, defined as: 1 – (#UMIs / #Counts). This statistic provides a per-cell measurement of the completeness with which unique molecules are sampled per cell and has the benefit of being adaptable across diverse cell types. Data were filtered to require RNA saturation > 0.35. Analyses utilized the Seurat R package, version 4.2.^63^ Using standardized methods implemented in the Seurat R package, counts and UMIs were normalized across cells, scaled per 10,000 bases, and converted to log scale using the ’NormalizeData’ function. These values were then converted to z-scores using the ’ScaleData’ command. Highly variable genes were selected using the ’FindVariableGenes’ function with a dispersion cutoff of 0.5. Principal components were calculated for these selected genes and projected onto all other genes using the ’RunPCA’ and ’ProjectPCA’ commands. Clusters of similar cells were identified using the Louvain method for community detection, and UMAP projections were calculated. CITE-seq data were CLR-normalized by lane, meaning raw count data are subset per lane, CLR normalization performed (as implemented in the Seurat R package, using margin = 1), using all QC-passing cells/lane. Normalization per-lane was performed to reduce batch effects.

### TCR Sequence Analysis

Raw sequence reads for gene expression and TCR enrichment were first processed using cellranger software, version 6.1.1 (10X Genomics). The raw clonotype calls produced by cellranger vdj were extracted from the comma-delimited outputs. Cells were demultiplexed and TCR calls were assigned to samples using custom software, made publicly available through the cellhashR package, with the GMM-Demux, demuxEM, and BFF algorithms.^60–62^ The clonotype data generated by cellranger were filtered to drop any cells where the TCR calls lacking a CDR3 sequence, the clonotype was not marked as full-length, or the clonotype lacked a called V, J, or constant gene. Rows with chimeric V/J/C combinations (i.e. TRBV / TRAC) were filtered, with the exception that segments consisting of a TRDV/TRAC or TRAV/TRDC were permitted. These chimeric segments were classified according to the constant region chain. Data was analyzed in R studio using Seurat package.^64^ All code is available on github (https://github.com/kaindylan/MR1T-Delayed-BCG/tree/main).

### TCR Group Clustering

TCR group clustering was performed by isolating V/J gene segments and CDR3 sequences from cells from the 10x datasets, along with post-hoc manual annotation of V/J gene segments and CDR3 sequences from the *Mtb* reactive clones. CoNGA was used to compute TRB TCR distances.^33,65^ Those distances were clustered by transforming via their first 50 kernel principal components,^66^ embedding in a k nearest neighbors’ graph (k = 323, or 10% of the graph size), and Leiden clustering^67^ using resolution parameter 1.1 and 10 iterations.

### Limited Dilution Cloning and Flow Confirmation of Ligand Identification

These results were published previously by Harriff *et al* and Kain *et al.*^31,35^ CBMC were stained with Propidium Iodide (Milteny Biotec), MR1-5-OP-RU tetramer (NIH tetramer core), and antibody cocktail of surface stains to sort out non-CD3+ cells (listed in Supplemental Table 8). Live, MR1/5-OP-RU tetramer positive cells were sorted and then cryopreserved. After thawing, the cells were plated in a 96-well plate in a limited dilution assay with 1.5x10^5^ irradiated PBMC (3000 cGray) and 3x10^4^ irradiated LCL (6000 cGray) along with αCD3 (30ng/ml) and IL-2 (2ng/ml) in RPMI 1640 supplemented with 10% heat inactivated human serum in 96 well round bottom plates. On day 5, the αCD3 was washed out by removal of half of the volume from the wells and replaced with additional IL-2 supplemented media. Media was changed for all wells every 2-3 days and replaced with IL-2 supplemented media. T cell “buttons” were evaluated for growth on Day 20 and selected buttons stained with the MR1-5-OP-RU tetramer, CD3 PeCy7 (clone SK7; Biolegend), CD4 FITC (clone SK3; Biolegend), CD8 APC Cy7 (clone SK1; Biolegend), and TRAV1-2 BV605 (clone IP26; Biolegend). T cell clones that remained MR1-5-OP-RU tetramer positive were re-expanded in T12.5 flasks with irradiated PBMC and LCL as well as RPMI 1640 media with 10% Human serum supplemented with αCD3 and IL-2 as above.^68^

### Statistics

Statistical analysis was done in R studio for all differential gene expression of single-cell gene expression data. Wilcoxon test for statistical significance of gene expression between different groups. For all analysis we considered a 2-log threshold change of more than 0.5 and a Bonferroni-corrected p-value of < 0.05 to be significant. For TCR cluster enrichment statistical significance was determined using a Fischer exact test with Holm-Bonferroni correction. For TCR pairing analysis, data was first analyzed in R studio and then diversity and pairing preference data was extracted and analyzed in Prism. Statistics were performed in Prism using a t-test for significance, with p < 0.05 being considered significant. Samples were all processed and analyzed in a blinded fashion with respect to BCG vaccine status.

### Data Availability

All R studio code used in the generation of this analysis is freely available and uploaded to github (https://github.com/kaindylan/MR1T-Delayed-BCG/tree/main). All raw data was deposited into the Sequence Read Archive database. The authors declare that the data supporting the findings of this study are available within the paper and its supplementary information files. The source data underlying the graphs in the paper can be found in the Supplementary Data.

## Author Contribution

DK, TJS, DML, DAL contributed to the conception and/or design of the work. EN, WH, MS, GW, NDP contributed to clinical activities. DML, DAL raised grants to fund the research. DK, GWM, GMS, KR, GB, EN, TJS, BB, DML, DAL substantially contributed to the acquisition, analysis, or interpretation of data and drafting of the manuscript. All authors substantially contributed to revising and critically reviewing the manuscript for important intellectual content. All authors approved the final version of this manuscript to be published and agree to be accountable for all aspects of the work.

## Acknowledgements

We would like to thank the participants who gave time and dedication to this health research as well as the SATVI field site clinical and laboratory teams (Libby Briel, Helen Veldtsman, Nondumiso Khomba, Bernadette Pienaar, Hadn Africa, Marcia Steyn, Ashley Veldsman, Humphrey Mulenga). We would also like to thank the BioMedical Research Institute (BMRI) Clinical Team and the Stellenbosch University Immunology Research Group laboratory team. The MR1 tetramer technology was developed jointly by Dr. James McCluskey, Dr. Jamie Rossjohn, and Dr. David Fairlie, and the material was produced by the NIH Tetramer Core Facility as permitted to be distributed by the University of Melbourne. We acknowledge the assistance of the Oregon Clinical & Translational Research Institute, which is supported by the National Center for Advancing Translational Sciences, National Institutes of Health, through Grant Award Number UL1TR002369. This project used the OHSU Flow Cytometry and Monoclonal Antibody Shared Resource Core Facility (RRID:SCR_009974).

