## Supplemental Figures for "BCG Vaccination at Birth Shapes the TCR Usage and Functional Profile of MR1T Cells at 9 Weeks of Age"

**Supplemental Figure 1: Representative FACS Sorting Strategy.**

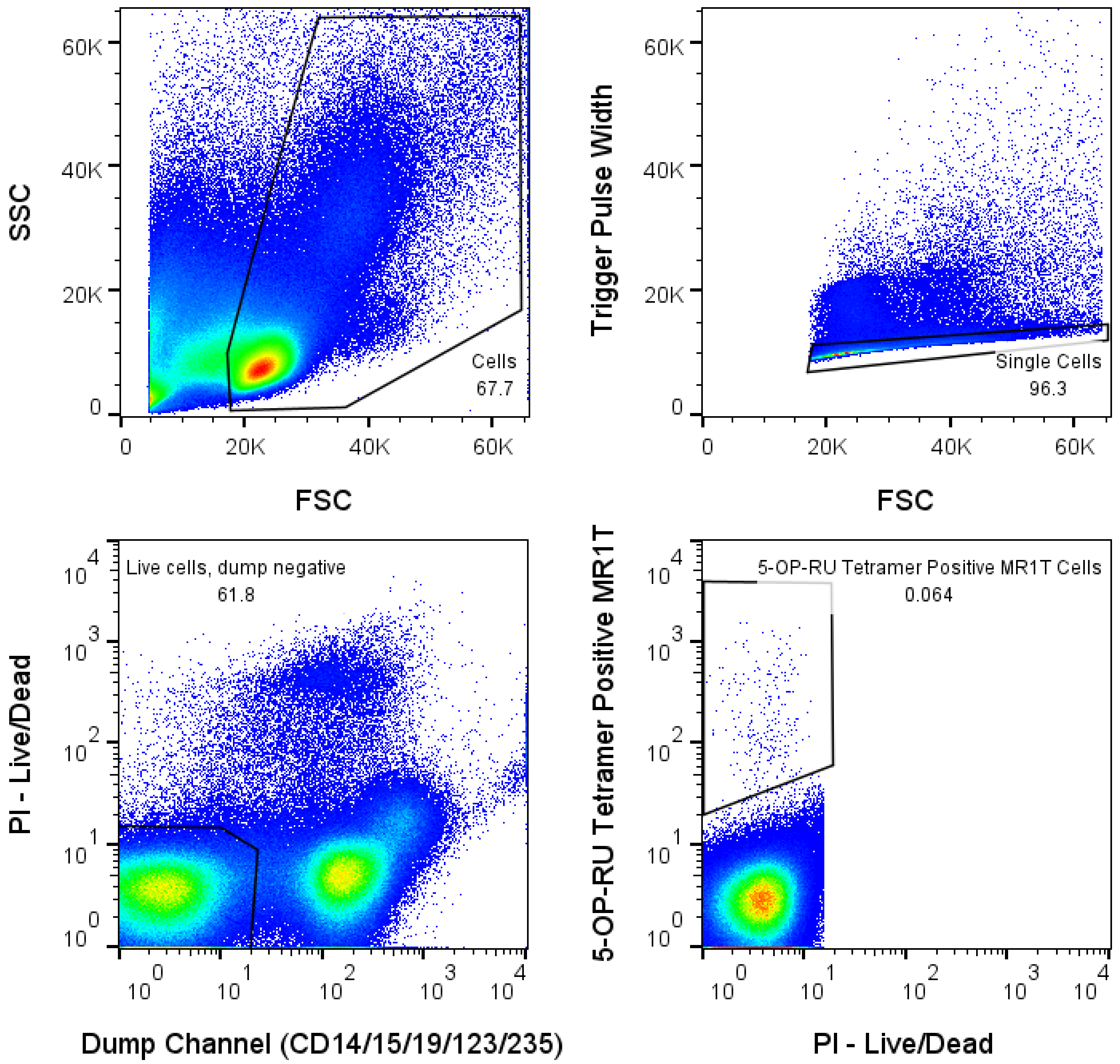

### Supplemental Figure 2: Phenotypic Features of MR1T cells.

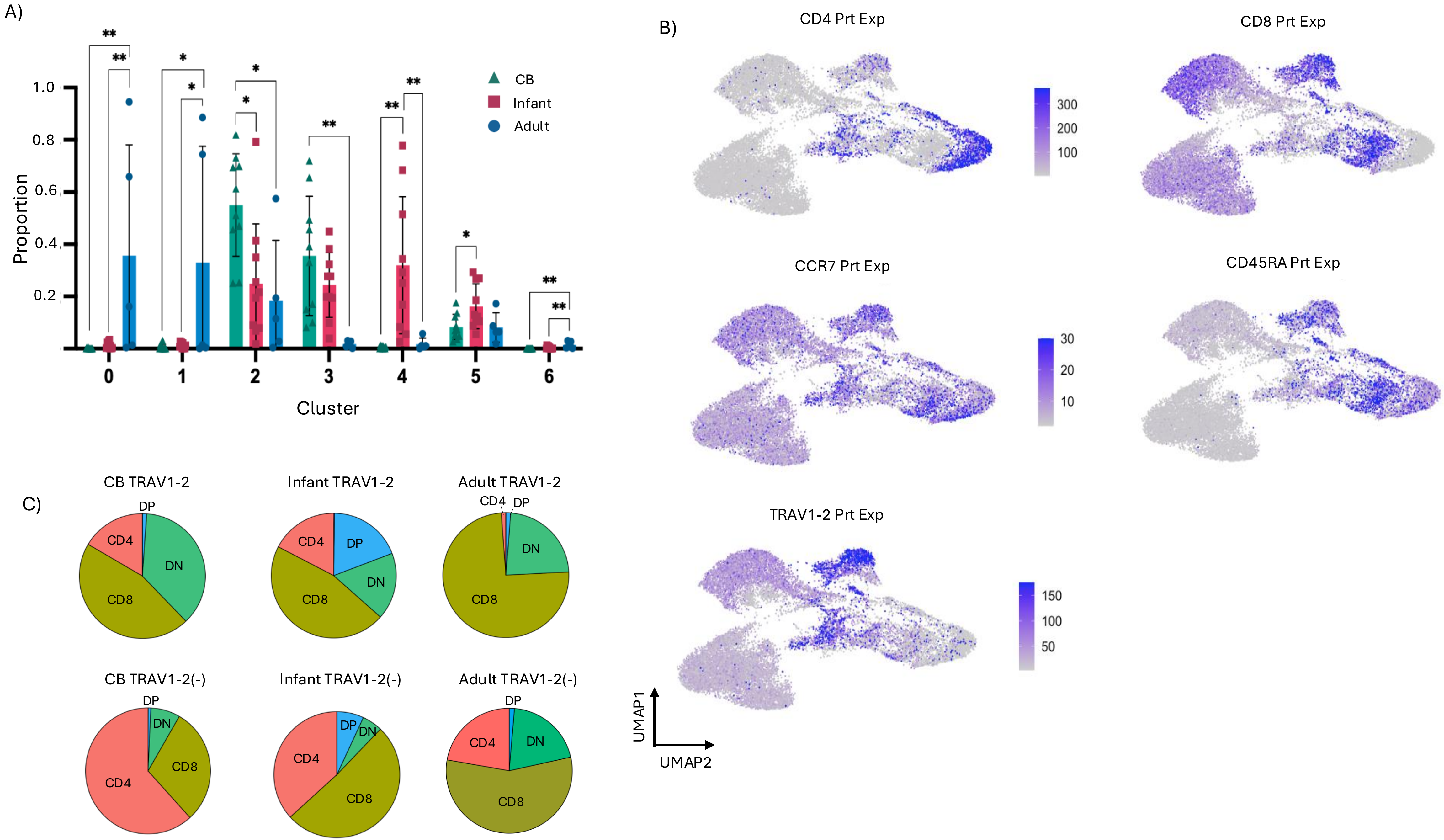

### Supplemental Figure 3: Age related changes in MR1T cells.

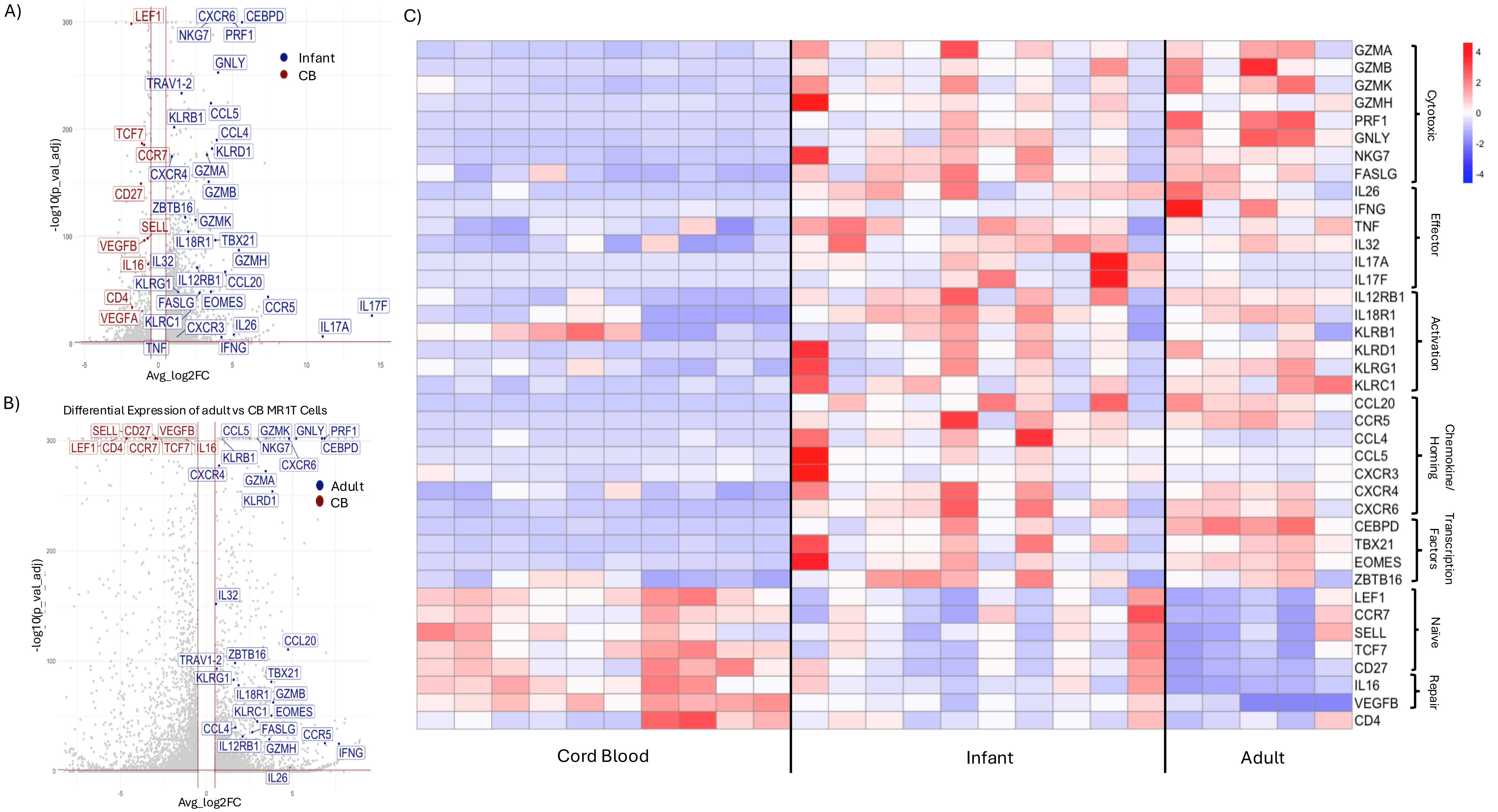

### Supplemental Figure 4: MR1T cell CDR3α length diversity.

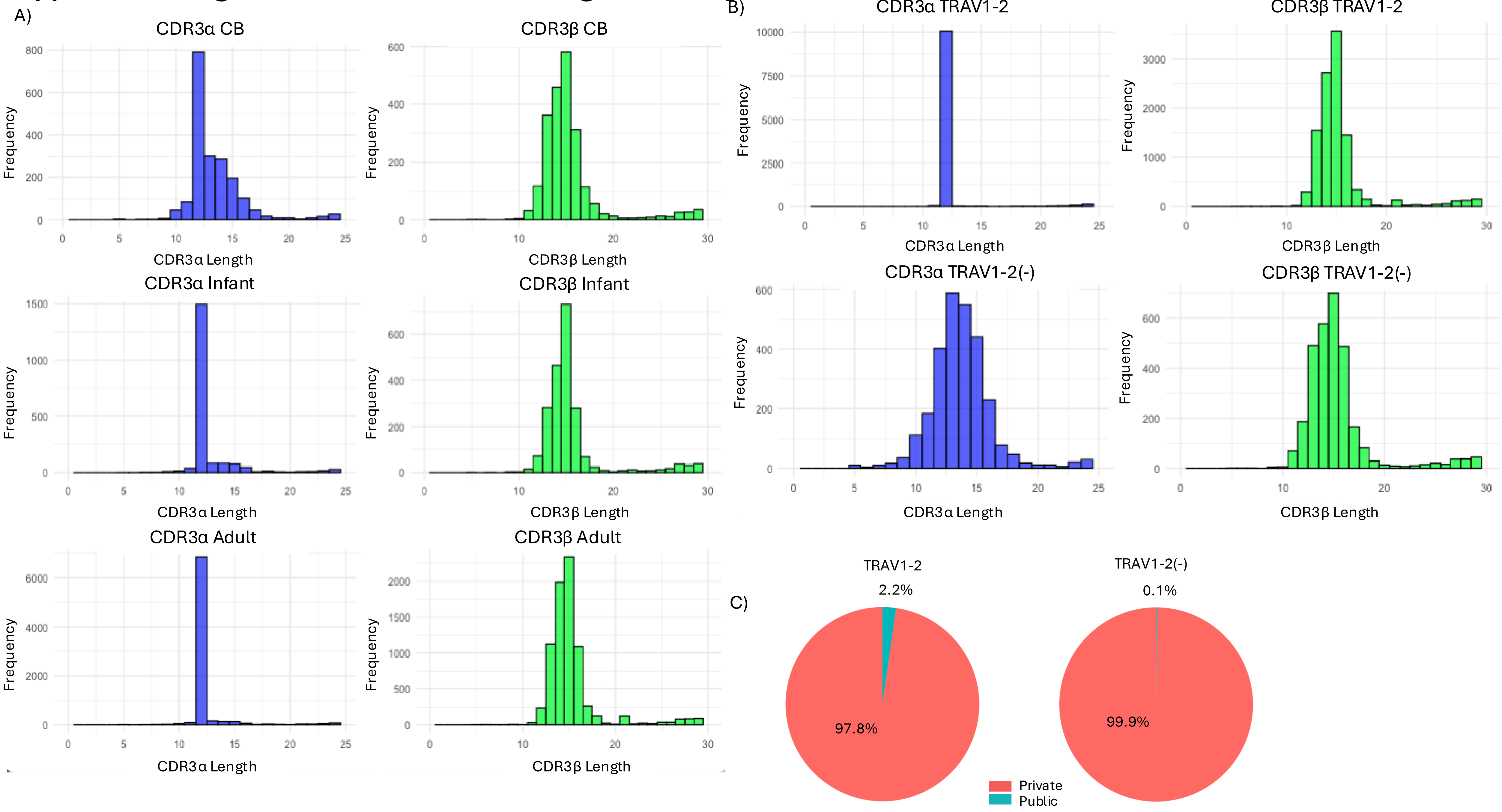

**Supplemental Figure 5: Gene expression in individual donors.**

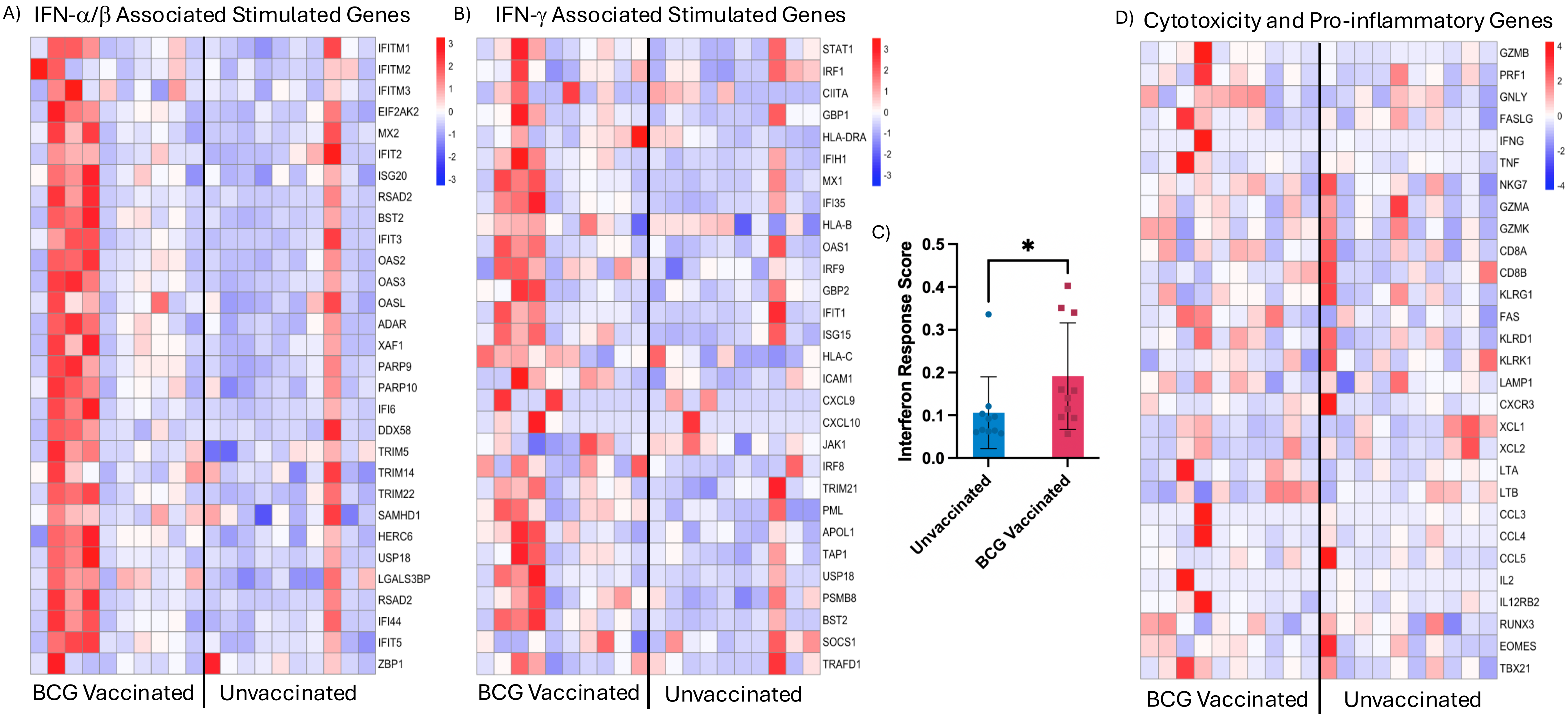

### Supplemental Figure 6: Cell surface protein and gene expression of infant MR1T cells.

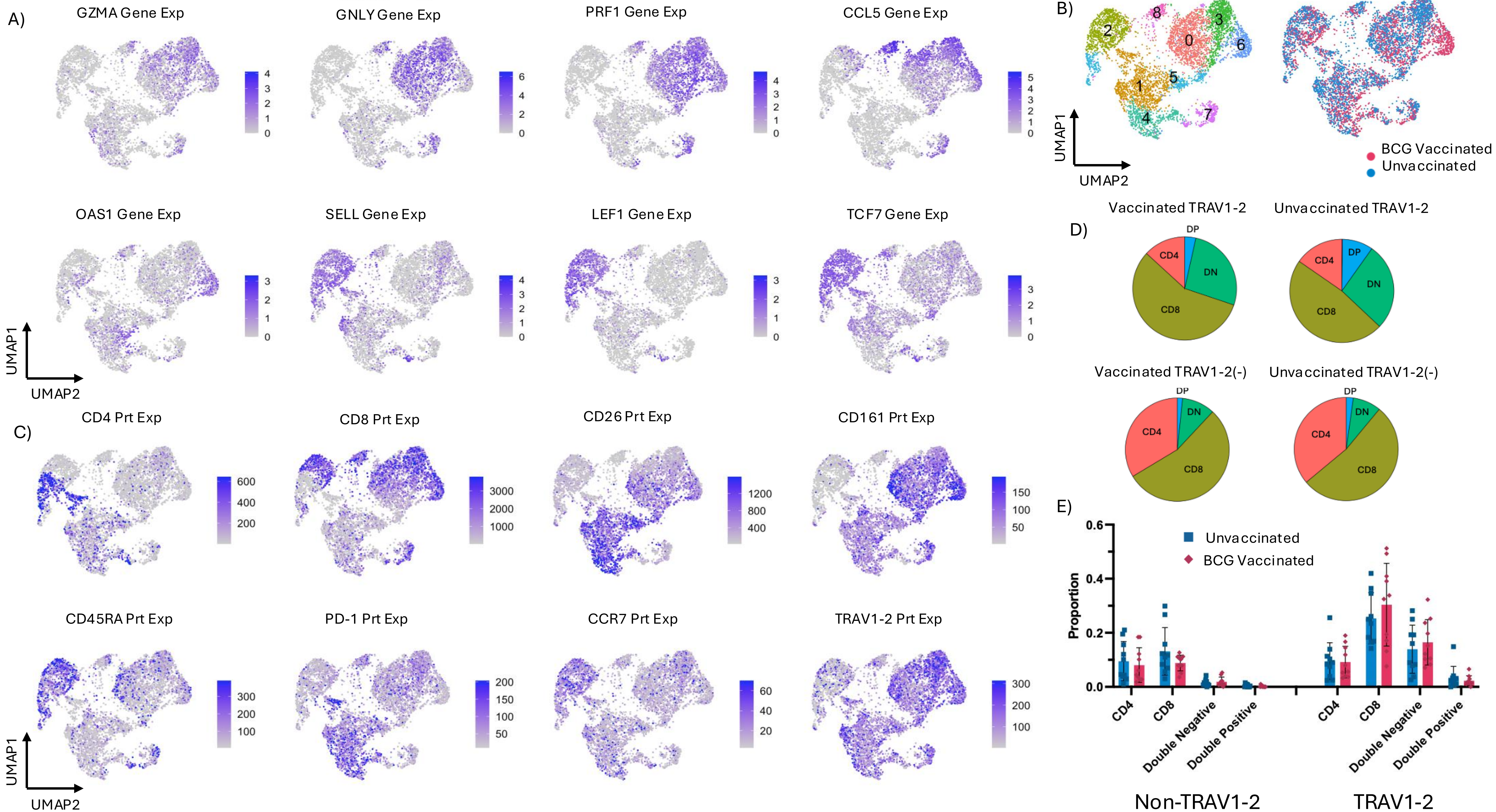

**Supplemental Figure 7: No differences in TRAV, TRAJ or TRBV usage with BCG Vaccination.**

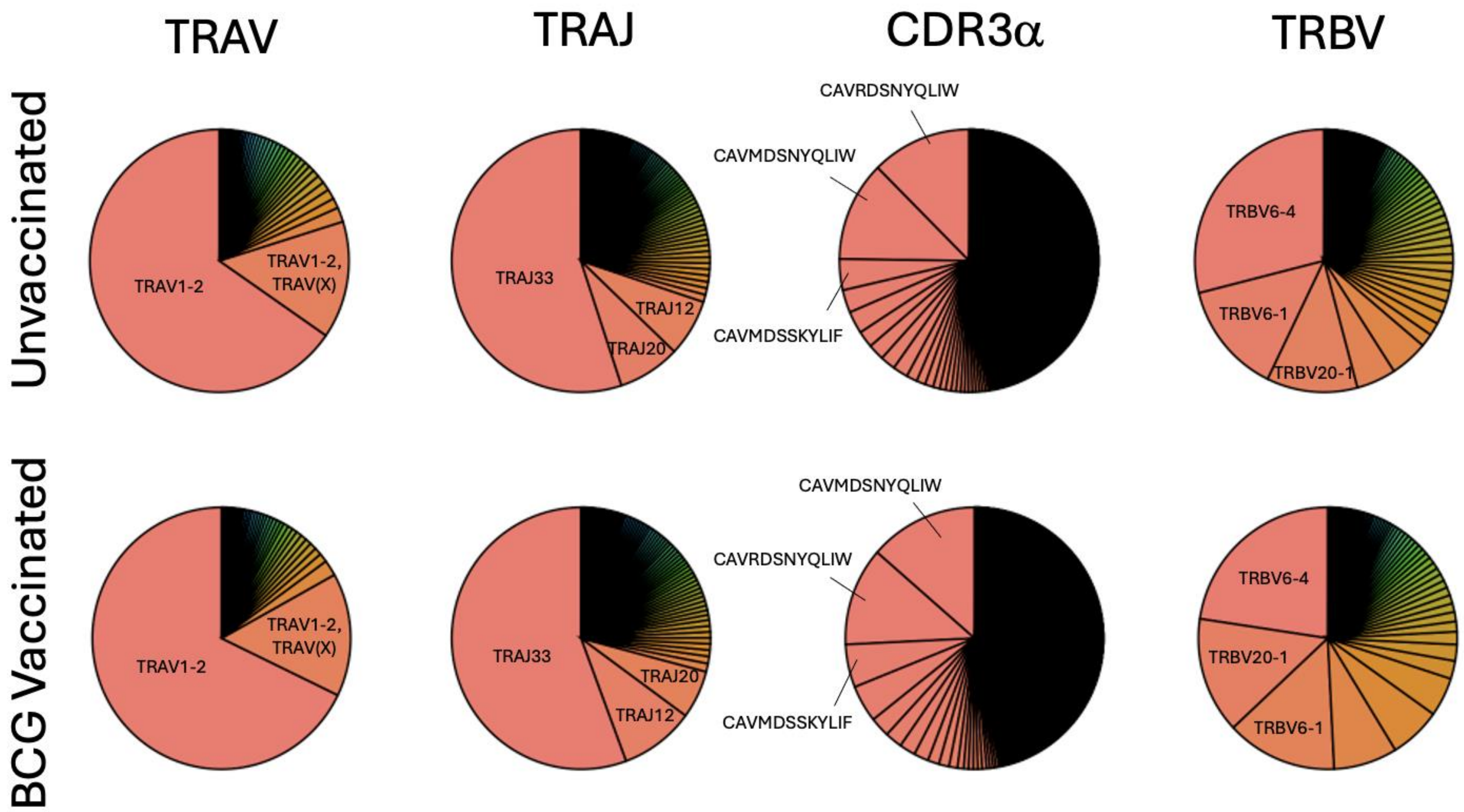
